# Absolute quantification of rare gene targets in limited samples using crude lysate and ddPCR

**DOI:** 10.1101/2024.03.27.586936

**Authors:** Charandeep Kaur, Stuart Adams, Catherine N Kibirige, Becca Asquith

## Abstract

**Background:** Accurate quantification of rare genes from limited clinical samples is crucial for research purposes but is technically challenging. In particular, nucleic acid extraction for quantification of gene targets may lead to target loss. Here, we report the development and validation of a novel crude lysate ddPCR assay for the absolute quantification of rare genes, TRECs in our case, from infrequent cells, that removes the need for DNA extraction, hence minimizing the target loss.

**Methods:** The analytical validation was performed on PBMCs extracted from the blood of healthy donors. Standard ddPCR was first optimized to detect TREC copies/cell and then applied to a crude lysate ddPCR assay. The assay was optimized by varying several steps. The optimised assay was directly compared to standard ddPCR and the performance of the assay quantified.

**Results:** The newly developed assay showed good agreement with the standard ddPCR assay in the range from 0.0003 to 0.01 TRECs/cell. The assay had a limit of quantification of <0.0003 TRECs/cell and a limit of detection of <0.0001 TRECs/cell; this performance is favourable compared to standard ddPCR. The intra-assay variation was low. This method can also be applied to fixed and permeabilized cells.

**Conclusions:** The newly developed crude lysate ddPCR assay for quantifying rare targets from limited samples has high accuracy, specificity, and reproducibility; additionally, it eliminates the need for DNA extraction for absolute quantification. The assay has the potential to be used for quantification of other trace targets from small samples.

## Introduction

Recent technological advances have transformed digital polymerase chain reaction (dPCR) from being an expensive technique with limited application, to a readily-accessible, widely-applied technology (1). With its exceptional analytical sensitivity and precision over quantitative PCR/ real-time PCR (qPCR/RT-PCR) (2–4), dPCR can be utilized for analysis of gene expression (1), epigenetics (2), copy number variation (3–5), rare mutations (6), linkage studies (7), and quantification of next-generation sequencing libraries (8).

In dPCR, a sample is divided into many small partitions, each of which undergoes PCR amplification. Poisson statistics are used to determine the absolute quantity of a target sequence in a sample based on the number of partitions in which the sequence is detected (9,10). This approach permits absolute quantification without the need of a standard curve, offering an advantage over qPCR (9).

Despite many advances in dPCR technology, its utility and performance still pose a challenge when the amount of starting material is limited. This problem is partially solved by droplet digital PCR (ddPCR); a form of dPCR that uses water-oil emulsion droplets to partition the sample (10). ddPCR can work with nM to pM concentrations of DNA (10). However, DNA extraction from ∼<1000 cells often results in the loss of the target during the extraction process. There is therefore a need for a high throughput, sensitive and accurate assay for the detection of rare targets from limited clinical samples.

Here we report a study to quantify T-Cell Receptor Excision Circles (TRECs), a rare gene target, in peripheral blood mononuclear cells (PBMCs) and sorted memory T cells using optimized crude lysate ddPCR. TRECs are small loops of extrachromosomal DNA formed in T cells during V(D)J recombination of the T cell receptor genes in the thymus. TRECs are not replicated during mitosis and so are diluted in the cell population with each round of cell division (11). TREC measurement is used to quantify thymic output and cell division history, and is implemented in newborn screening programs for Severe Combined Immunodeficiency (12). Different subpopulations of T cells have different concentrations of TRECs depending on the number of cell divisions they have undergone since thymic production. Consequently, the naïve T cell subpopulation (Tn) has markedly higher TREC concentrations than memory T cell subpopulations such as effector memory T cells (Tem), central memory T cells (Tcm) and T stem cell memory (Tscm) (13). Quantification of TRECs in low-frequency memory T subpopulations e.g. T stem cell memory cells (Tscm) (which constitute 2-4% of the total CD4+ and CD8+ T cell population in blood, (13,14) or antigen-specific T cells (in absence of acute infection, one cell within 100 to 10^5^ T cells (15), are problematic for two reasons: the target is rare and the starting cell numbers low. We, therefore, sought to develop a sensitive and accurate ddPCR method directly from crude lysate.

## Material and Methods

### Peripheral blood samples and monocyte cell line

Peripheral blood samples (60-70ml) from healthy donors were collected in Sodium Heparin tubes (BD Vacutainer) and processed within 3h. PBMCs were isolated via density gradient centrifugation over Histopaque®Hybri-max (Sigma-Aldrich). The PBMC layer was collected, washed with PBS, and cryopreserved in 10% DMSO (Dimethyl Sulfoxide) (Merck) in FCS (Fetal Calf Serum) (heat-inactivated, Gibco).

When required, cryopreserved PBMCs were revived in RPMI-1640 media (Sigma) supplemented with 10% FCS. Cells were counted by acridine orange/propidium iodide staining and analyzed using Luna dual Fluorescence cell counter (Logos Biosystems, South Korea). A fraction of these cells was used to prepare standard curve dilutions while the rest of the cells were used for DNA extraction (see “DNA extraction”, Materials and Methods). A monocyte cell line (U-937)/macrophage DNA was used as a negative control.

### Sorting of CD8^+^ T memory subpopulations

For the direct comparison of the newly developed crude lysate ddPCR and standard ddPCR, CD8+T memory samples (rather than cryopreserved PBMCs or T cells diluted with macrophages) were used. For this, PBMCs were isolated from the blood of healthy volunteer and enriched for CD3^+^ Tcells using EasySep™ human T-cell isolation kit (Stemcell Technologies, Vancouver,Canada) as per manufacturer’s instructions. Enriched CD3^+^ T cells were stained with the following fluorochrome-attached antibodies: CD8 (BV711, Biolegend), CD45RA (ECD, Beckman Coulter), CD27 (Qdot605, ThermoFisher), CCR7 (FITC, BD BioScience) and CD95 (Pe-Cy5, Biolegend) antibodies. Stained cells were sorted into different memory subpopulations using a BD FACSAria III-U (BD Biosciences).

### Cell fixation and permeabilization

To investigate whether cell lysate buffer works on fixed and permeabilized cells, 2*10^6^ PBMCs were fixed using 200ul of fixation buffer (Invitrogen) for 30mins at room temperature. Fixed cells were washed with 200ul of 1% permeabilization buffer (Invitrogen) and then permeabilized in 200ul of 1% permeabilization buffer for 20 mins.

### DNA Extraction

Genomic DNA from PBMCs and sorted memory CD8+ T cell subpopulations was extracted using DNeasy Blood and Tissue Kit (Qiagen) as per manufacturer’s instructions. DNA was eluted in 60ul of nuclease free water (NFW) and stored at –20°C. DNA concentration was measured fluorometrically by Qubit 3.0 (Invitrogen).

### Cell lysate preparation

Five different approaches to preparing the cell lysate were compared:

1. Thermal lysis in NFW. 1000-10,000 cells were resuspended in 10ul of NFW and boiled for 5 mins at 99°C on water bath. 10ul of ProteinaseK (10mg/ml) was added and cells were heated at 57° C for 30 min, followed by 99°C for 30 mins. Next, cells were rapidly cooled on ice and centrifuged for 2 min at full speed. The supernatant was used for ddPCR.
2. Sonication. For sonication lysis protocol, 1000-10,000 cells were resuspended in 50µl of NFW and sonicated for 1 cycle of 30sec with Peak Power= 105, Duty factor= 10 and cycle/burst=200.
3. Lysis buffer from DNeasy Blood & Tissue Kit (Qiagen). Only lysis reagents were used as per manufacturer’s instructions.
4. Lysis in Buffer 1. Lysis reagents from the Ambion Cell to-Ct® kit (ThermoFisher) were used. Cell lysate was prepared according to manufacturer’s instructions, omitting the DNase DNA degradation step.
5. Lysis in Buffer 2. Lysis reagents from the SuperScript™ IV CellsDirect™ cDNA Synthesis Kit (ThermoFisher) were used. Cell lysate was prepared according to manufacturer’s instructions, omitting the DNase DNA degradation step.

Cell lysate prepared using the methods above was used in the ddPCR reaction mixture.

### Cell lysate viscosity breakdown step

A cell lysate viscosity breakdown step was investigated to completely lyse the cells and to reduce the viscosity of lysate. The cell lysate was heated on a thermocycler at 65°C for 1min, 96°C for 2min, 65°C for 4min, 96°C for 1 min, 65°C for 1 min and 96°C for 30 sec. Tubes were spun down for 1 min and cooled at room temperature before using in ddPCR reaction mixture.

### ddPCR reaction mixture preparation

All the preparation steps and reaction set up were performed in a dedicated pre-PCR room and in a dedicated PCR hood. Two probes were used: a TRECs probe labelled with FAM and a housekeeping gene RPP30 (ribonuclease P/MRP subunit p30) probe labelled with HEX. The sequences of primers and probe used for TRECs are: Forward primer 5’-CAC ATC CCT TTC AAC CAT GCT-3’; Reverse primer 5’-GCC AGC TGC AGG GTT TAG G-3’, Probe Sequence: 5’-ACA CCT CTG GTT TTT GTA AAG GTG CCC ACT-3’ with FAM and BQ-1 quencher (16). For RPP30, we used the primer probe mix for copy number detection from BioRad.

A 22ul reaction mixture was prepared comprising 11ul 2×ddPCR Supermix for probes (no dUTP) (Bio-Rad, CA, USA), 0.55 ul of TRECs primers (36uM stock) and probes (10uM) each, 1.1 ul of RPP30 copy number detection (Bio-Rad), 1ul of HindIII-HF restriction enzyme (NEB, UK). The remaining 7.8ul was adjusted with NFW and extracted DNA for standard ddPCR. Or, for crude lysate ddPCR, 7.8ul of cell lysate was used. For buffer 1, seven replicates were made and for buffer 2, four replicates were made. During data analysis, these replicates were summed so that a total of ∼140,000 droplets/sample were screened for buffer 1 and 80,000 droplets/sample for buffer 2.

### ddPCR droplet generation and thermocycler protocol

From the 22ul of ddPCR reaction mixture, 20 μl was loaded into a droplet cartridge (Bio-Rad, USA) in sample loading wells. 70ul of droplet oil was loaded in the droplet oil wells. The cartridge was covered with a gasket and placed in the droplet generator (Bio-Rad #186-3002). 40ul of droplets generated in the cartridge were transferred to a 96-well plate (Bio-Rad, USA) using a multichannel pipette. The plate was sealed using foil in the plate sealer (Bio-Rad, USA). The plate was put on the thermocycler (Bio-Rad, USA) under the following conditions: 10 min hold at 95 °C, 45 cycles of 94 °C for 30 s then 59 °C for 60 s, next step of 98 °C for 10 min and final hold at 12 °C. Amplified droplets were then read in the QX200 Droplet Reader (Bio-Rad, USA). The droplet reader was turned on for 30 mins before reading the plate.

### Droplet volume size estimated by optical microscopy

The average droplet volume was compared for standard and crude lysate ddPCR. Droplet generation and acquisition of the optical microscopy images were performed on the same day. Four wells were randomly selected for both of the ddPCR methods from different cartridges. 80-100 droplets/well were measured to get approximately 300 droplets in total. The droplets were transferred to μ-Slide VI flat uncoated microscopic chamber (IBIDI Germany). The slide was held at an angle for a few seconds to obtain a uniform monolayer of droplets. An optical microscope (Leica SP8 confocal microscope) with a digital camera was used to image the droplets. Images were recorded under uniform illumination in a bright field imaging mode, with 200x magnification. The 2-D images were analyzed in ImageJ (2.9.0) following an existing procedure described by (17).

### Assay linearity

The linearity and accuracy of the standard ddPCR and crude lysate ddPCR were assessed separately. These assessments ensure that the assay output is directly proportional to the input concentration of the target molecule and establish the assay’s quantitative reliability. For standard ddPCR assay, a standard curve of five serial dilutions (in 3 replicates) was made using CD3+ T cell DNA diluted with macrophage DNA (macrophages are negative for TRECs).

For crude lysate ddPCR using buffer 1 and buffer 2, 100-10,000 cells (depending upon number of TRECs copies/cell) were used to make two-fold serial dilutions in PBS covering a range of 2-64 TRECs copies/dilution. Serially diluted cells were then lysed using either buffer 1 or buffer 2 (details under heading Cell lysate preparation in methods). These linearity assessment experiments for lysis buffer 1 or buffer 2 were performed on two separate days using distinct PBMC samples, each with varying TREC copies per cell. The housekeeping gene RPP30 served as an internal control for quantifying cell numbers.

### Standard ddPCR assay accuracy relative to qPCR

For standard ddPCR, we assessed the accuracy of the assay by comparing its estimates of TREC content with those obtained by a well-established qPCR method developed by Dr Stuart Adam, from Great Ormond Street Hospital, London. We exchanged DNA samples and independently quantified TRECs copies/cell.

### Crude lysate ddPCR assay accuracy relative to standard ddPCR

The accuracy of the crude lysate ddPCR assay was evaluated on sorted memory T cell populations by comparing its estimates of TREC content with those obtained using the standard ddPCR.

### Limit of Blank

Limit of Blank (LOB) was calculated using the TRECs copy number concentration that was found in a total of 22 non template controls (NTCs) and 22 U-937 monocyte cell line lysate controls (CLCs) reactions using the formula (18):

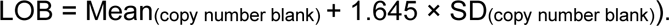

### Limit of Detection, and Limit of Quantification

A standard curve made from PBMCs diluted with the U-937 monocyte cell line was used to estimate the limit of detection (LOD) and limit of quantification (LOQ) of crude lysate ddPCR assay. A total of 4 dilutions were made with TRECs concentration ranging from

0.01 - 0.0002 TRECs/cell, ran in triplicates. The LOD and LOQ were calculated as previously described (18). Specifically, the LOD was the lowest copy number concentration that could be distinguished from the LOB with 95% certainty, calculated by the equation (18):

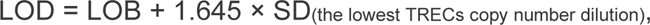

While the LOQ was defined as the lowest TRECs copies that could be detected with a coefficient of variation (CV) ≤35% (18).

### Repeatability of the ddPCR assay

To assess the repeatability (intra-assay variation) of the crude lysate ddPCR assay TREC copies/cell were measured in PBMCs derived from 4 different donors and ran in triplicates. CV% was calculated to assess the repeatability.

#### Data analysis

The ddPCR data were analysed in QuantaSoft analysis software version 1.7 (Bio-Rad). In each experiment, 3-8 NTCs and CLCs reactions were used to define the threshold that differentiates the positive droplets from negative droplets. The reactions with >8000 accepted droplets/well were used for the analysis. The copy number concentration (copies/ul) was calculated using the formula:

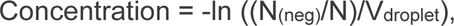

where N_(neg)_ is number of negative droplets, N is the total droplet and V_droplet_ is the volume of a droplet. We had previously determined V_droplet_ to be 0.70ul by optical microscopy. For crude lysate ddPCR, TRECs/cell were calculated by adding the replicates, seven in case of Buffer 1 and four for Buffer 2. The number of replicates depended on the total volume of the lysate. Linear regression, Spearman’s correlation coefficient, Bland Altman analysis and t-test were performed in GraphPad Prism version 9.4.1.

## Results

### OPTIMIZATION AND VALIDATION OF STANDARD ddPCR USING EXTRACTED DNA

In order to have an assay to compare the novel high throughput crude lysate ddPCR assay we first optimized and validated an in-house standard ddPCR assay to quantify TREC copies using extracted DNA from PBMCs. Annealing temperatures for TRECs and RPP30 primers and probes were optimized by performing a thermal gradient of 58°C - 62°C degrees. An optimum annealing temperature for both TRECs and RPP30 was determined to be 59°C as this provides the best separation between positive and negative droplets with minimum rain in between them, for both the channels (Fig 1a).

**Fig 1:**
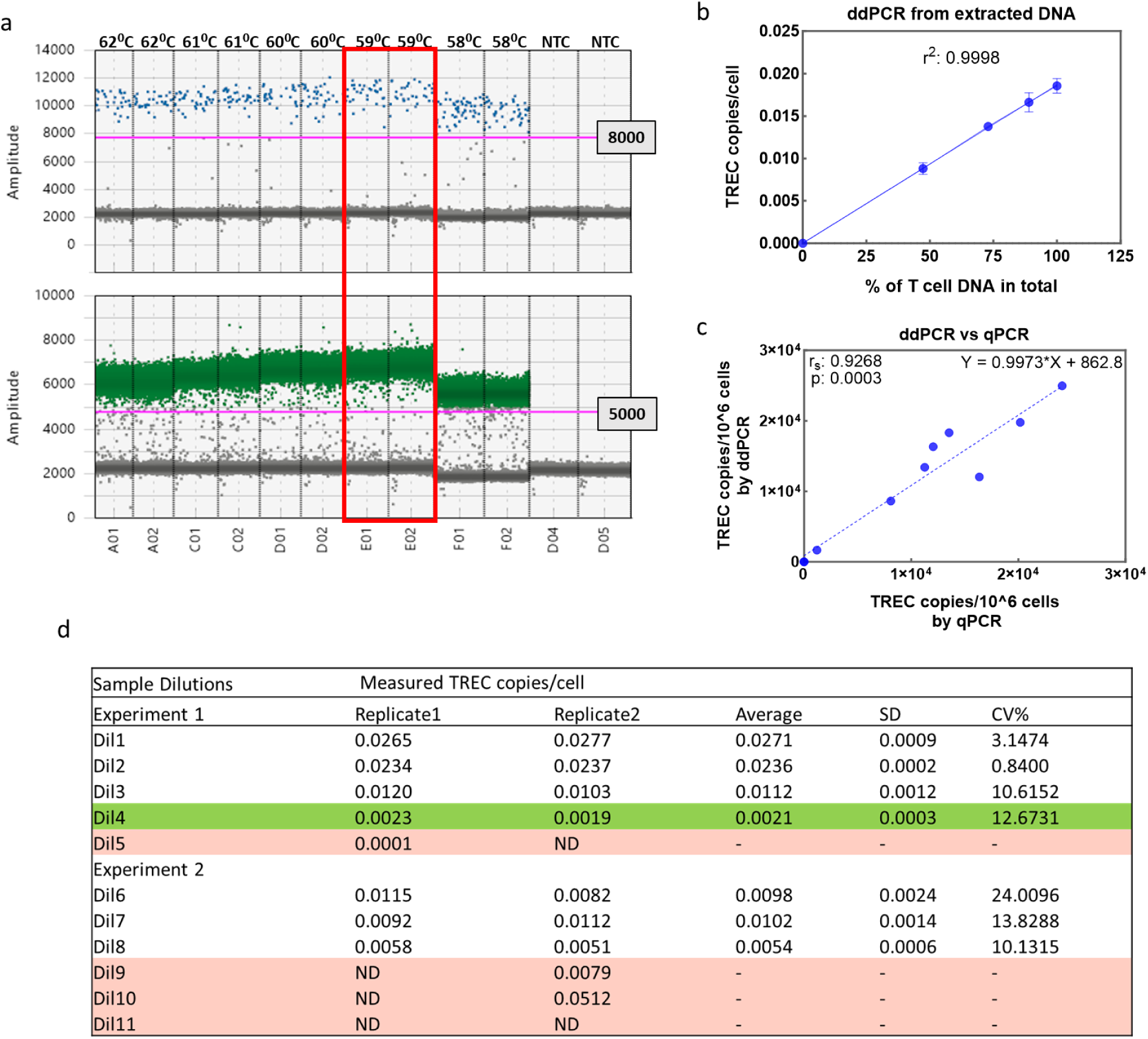
Optimization and validation of standard ddPCR. (a) Gradient temperature experiment was performed on DNA samples isolated from the blood of a healthy individual and run in duplicates. The upper panel shows a 1-D dot plot displaying the results for the amplitude of TRECs-positive droplets (blue), while the bottom panel presents the results for RPP30, a housekeeping gene(green). The pink horizontal line represents the threshold for positive droplets, and the red squares mark the optimal temperature selected based on the best separation and minimum rain. (b) A standard curve was made by diluting T-cell DNA with macrophage DNA to assess the linearity of the assay. Samples were run in duplicate, the coefficient of determination r_2_ calculated by linear regression. (c) To measure accuracy, TREC copies/million cells were measured in n=10 DNA samples using both qPCR and ddPCR. (d) Dilution experiments for LOB, LOD and LOQ estimation. Green row shows the lowest dilution with TRECs detected in both the replicates. Pink rows highlight the dilutions with ND (not detected) in one or both the replicates. SD: Standard Deviation; CV: Coefficient of variation

Next, the linearity of this assay was investigated using serial dilutions of TRECs containing T cell DNA with TREC negative macrophage DNA. TREC copies/cell measured by ddPCR was plotted against percentage of T cell DNA in the reaction mixture and a straight line fitted by linear regression (Fig 1b). The best fit straight line had a coefficient of determination (r^2^) of 0.99, indicating good linearity between the TRECs copies/cell and the concentration of T cell DNA present in the sample.

Accuracy was assessed by comparing TREC copies/cell quantified from diluted PBMC samples by the standard ddPCR assay with the qPCR method. We observed a Spearman coefficient (r_s_) 0.93, with p: 0.0003 between the results obtained by the standard ddPCR and the qPCR assays (n=10) (Fig 1c). We found the equation of the straight line to be Y = 0.9973*X + 862.8, indicating excellent agreement between the methods over the range studied.

### PERFORMANCE OF THE STANDARD ASSAY

LOB, LOD and LOQ of standard ddPCR was assessed (Fig 1d). LOB for this assay is zero since we did not detect a single droplet for TRECs in NTCs. TREC copies/cell were detected in the dilutions having ≥ 0.002 TREC copies/cell, however below this concentration we did not detected TRECs in either one or both the replicates (highlighted in pink in Fig1d). Hence, we selected the Dil4 with 0.002 TRECs/cell (highlighted in green in Fig 1d) to estimate the LOD, calculated to be 0.0004 TRECs/cell (LOD: 0+1.645*0.0003). Further, this is the lowest dilution having 100% hit rate for probit analysis and CV%<35, Therefore, we estimate the LOQ to be 0.002 TRECs/cell (coefficient of variation (CV) 12.7%).

These findings confirmed that TREC copies measured by the standard ddPCR method is accurate and comparable with the well-established qPCR method. Henceforth, we use this standard ddPCR assay as a benchmark for our novel crude lysate ddPCR assay.

### OPTIMIZATION AND VALIDATION OF CRUDE LYSATE ddPCR ASSAY

We investigated different methods for preparing cell lysates. Firstly, sonication and thermal lysis were tested. Both of these methods suffered from a low rate of DNA recovery (supplementary, Fig S1). Next, lysis buffer from the DNeasy Blood & Tissue Kit (Qiagen) was tested and resulted in negligible droplet count presumably due to detergents or similar that burst the ddPCR droplets. Lysis Buffer 1 and Buffer 2 including the viscosity breakdown step were much more successful and taken forward for further analysis (Fig S1).

### IMPACT OF VISCOSITY BREAKDOWN PROTOCOL

To perform crude lysate ddPCR, we added an innovative step to make the cell lysates more compatible with ddPCR, called the viscosity breakdown step. The presence of intact oligonucleotides in crude cellular lysates elevates lysate viscosity, posing challenges for droplet formation and target amplification (19). To assess the impact of a viscosity breakdown (VB) protocol applied to lysed cells prior to droplet formation, we performed crude lysate ddPCR processed with and without the VB protocol (n=5) and compared the results with those obtained with standard ddPCR Fig 2a. We found higher levels of TREC copies/cell in the samples processed without the VB protocol (mean=0.046 TRECs/cell) compared to standard ddPCR (mean=0.023) (Wilcoxon test, p=0.06), while no significant difference was observed between VB protocol (mean 0.022) and standard ddPCR (mean=0.023) (Wilcoxon test p=0.44) (Fig 2a). These results suggest that the viscosity of the cell lysate may interfere with the amplification and/or incomplete cell lysis may be responsible for the higher TRECs levels observed in samples without the VB protocol. Consequently, we consistently utilized the VB protocol in conducting crude lysate ddPCR experiments.

**Figure 2:**
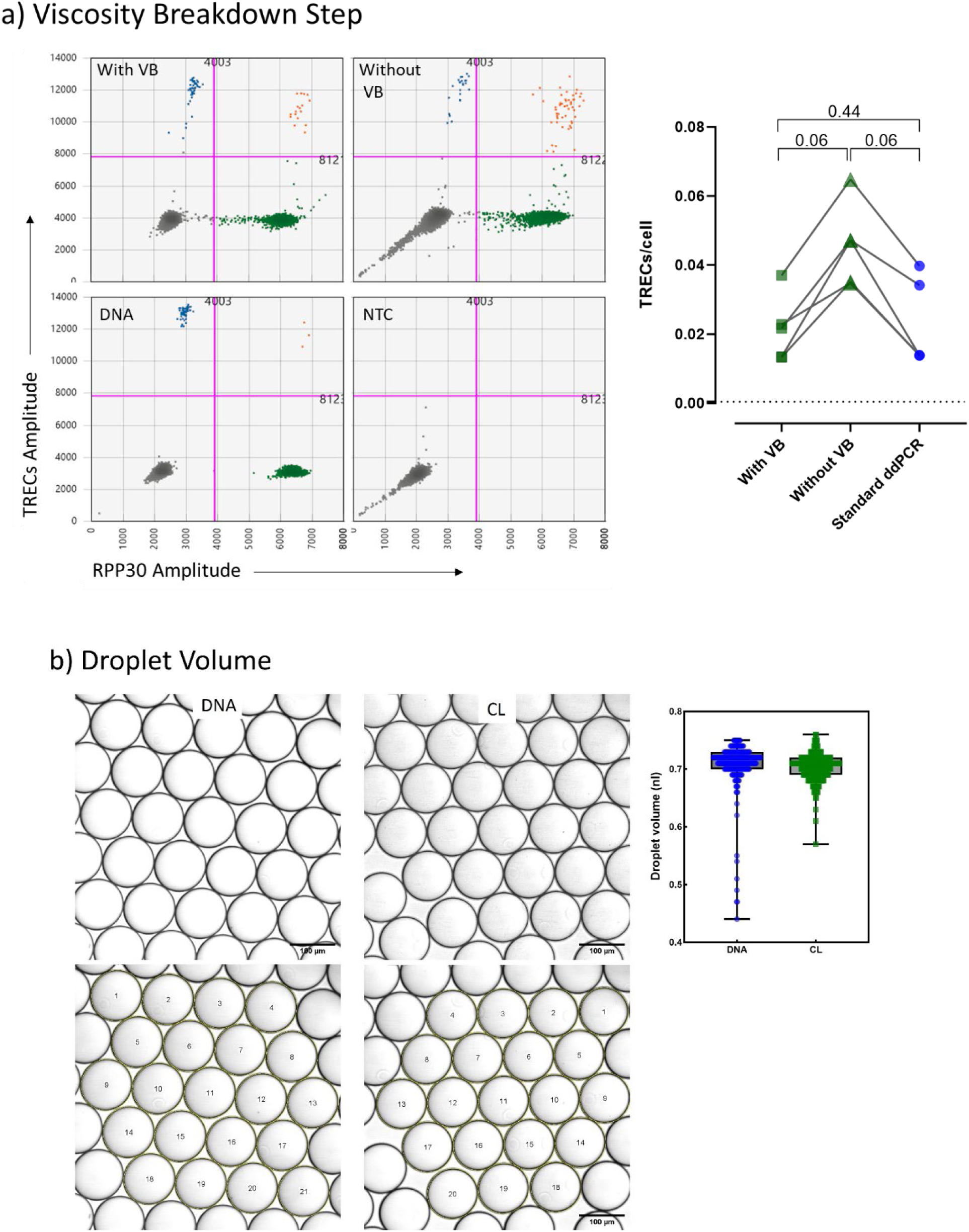
Importance of viscosity breakdown step and Droplet Volume measurement in crude lysate ddPCR. (a) 2-D dot plots show results of crude lysate ddPCR performed on PBMCs (one representative plot from 5 healthy individuals). Additionally, standard ddPCR (bottom left) and NTC (bottom right) were included as controls in this experiment. Blue dots are TRECs single positive, green dots are RPP30 single positive, Orange dots are TREC RPP30 double positive and grey dots are double negative. The pink line represents the threshold. Each dot represents a droplet in ddPCR. On the right hand -side, pairwise comparisons were made between the TRECs copies/cell measured by crude lysate ddPCR with VB (green squares) and without VB (green triangles) and the standard ddPCR (blue dots). Each symbol represents a donor, and pairwise comparison is shown by connecting black lines. Stats: Wilcoxon test. (b) Optical microscopy images are presented to measure droplet volume. On the left-side, a monolayer of droplets was generated using extracted DNA and crude lysate (CL) observed under a Leica SP8 Confocal (Germany) optical microscope. Bottom row shows the images after processing using ImageJ software which involved thresholding and analyzing particle size determination. On the right hand-side, the box-whiskers plot shows the median droplet size of droplets containing extracted DNA (in blue) and CL (in green), each circle represents one droplet.

### DROPLET VOLUME

We were concerned that the cell lysate buffers may affect the droplet volume. We therefore examined, by optical microscope, approx. 300 droplets from 4-5 different wells generated using crude lysate and compared this with droplets generated using extracted DNA. To prevent any potential alteration in droplet size during microscopic examination, the images were captured within 30 minutes of droplet generation.

A representative image of a droplet layer is provided in Fig 2b (top row) together with the image obtained after the image analysis in Fig 2b (bottom row). The average droplet volume generated using DNA was 0.7096nL (SD 0.037; 95%CI 0.7054-0.7138) and using CL was 0.7034nL (SD 0.022; 95%CI 0.7010-0.7058) (Fig 2b, right side), both of which are much smaller than the value of 0.85nL used in QuantaSoft version 1.0.596 to calculate copy number concentration.

Therefore, going forward, we used 0.70nL as the droplet volume in our calculations for both ddPCR using CL and DNA.

#### Assay linearity and accuracy

The linearity of the crude lysate ddPCR assay, prepared using both buffer 1 and buffer 2 was evaluated and compared with the standard ddPCR. TREC copies/cells were estimated from the standard curve dilutions covering a range of 2-64 TREC copies/dilution. For assay with Buffer 1, a strong linear relationship was observed between the number of cells/reaction (estimated from RPP30) and TREC copies, with an r² value of 0.95 (p < 0.001). (Fig 3a). However, measuring the accuracy by comparing the input TREC copies measured by the standard ddPCR with the output TREC copies measured by crude lysate ddPCR using Buffer 1 resulted in a linear regression equation of Y = 1.513X + 0.5430 (Fig 3c), showing that each unit increase in TREC values by standard ddPCR corresponds to a 1.513 unit increase in values from crude lysate ddPCR using Buffer 1. The gradient being >1 and the positive intercept (0.5430), suggesting a systematic overestimation.

**Figure 3:**
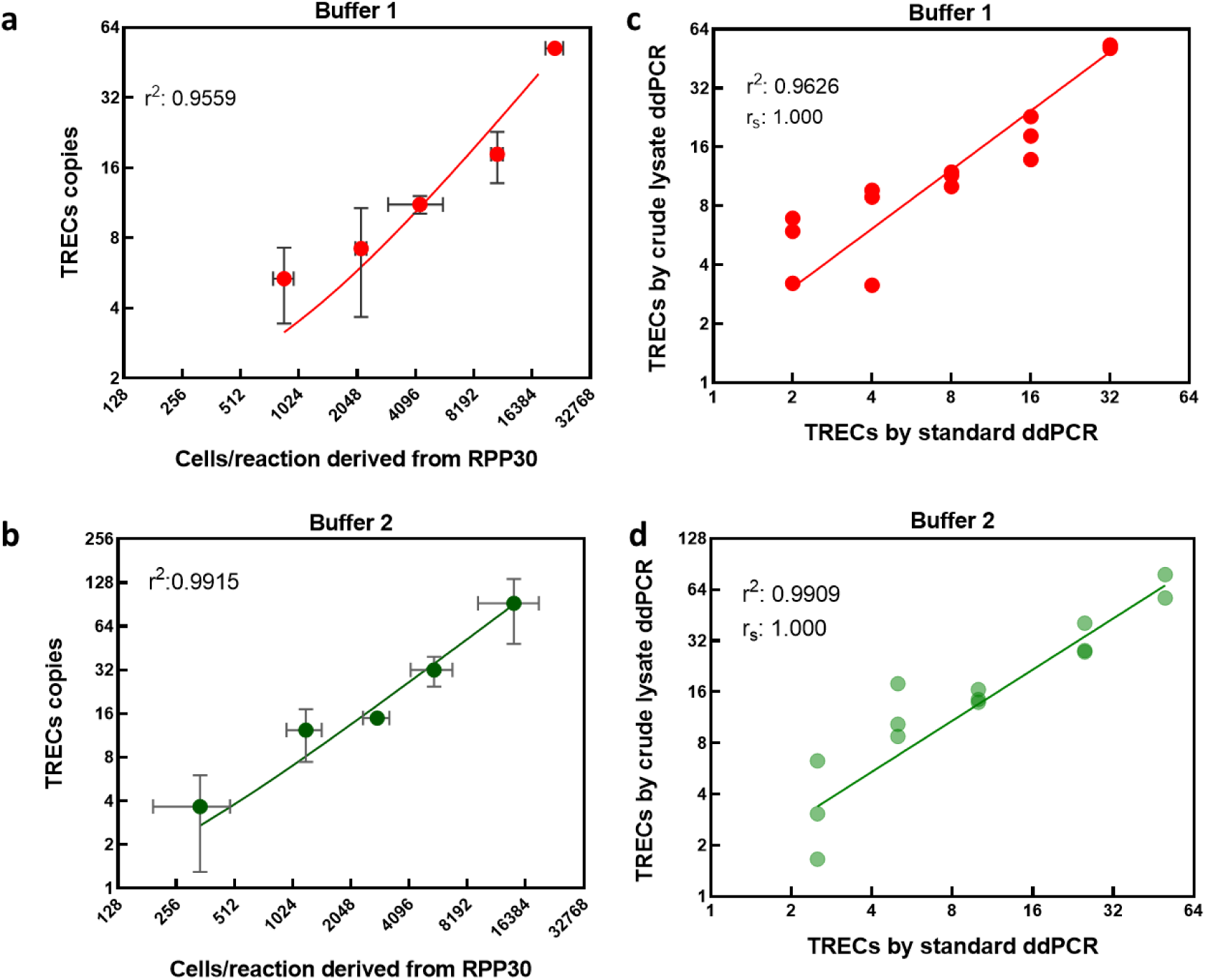
Linearity and accuracy assessment of crude lysate ddPCR using lysis buffer 1 and 2. (a-b) Linear regression analysis performed on the standard curve made by diluting PBMCs in PBS and then preparing crude lysate using lysis Buffer 1 (in Red) or Buffer 2 (in green). The PBMC samples used for Buffer 1 and Buffer 2 experiments were distinct, with varying TREC concentrations. (c-d) The accuracy of the assay was estimated by comparing the output TRECs concentration measured by crude lysate ddPCR using either lysis Buffer 1 (in red) or buffer 2 (in green) with input TRECs measured using extracted DNA by standard ddPCR. Linear regression equation for Buffer 1 is Y = 1.513X + 0.5430 and for Buffer 2 is Y = 1.298X + 0.6991. Spearman coefficient r_s_ =1 for both assays.

Assay with Buffer 2 also demonstrated good linearity between the number of cells/reaction and TREC copies, (r² value >0.99, p < 0.001) (Fig 3b). Further comparing the input TRECs estimated by standard ddPCR against the output TRECs estimated by the crude ddPCR, the linear regression equation (Y = 1.298X + 0.6991) indicates that each unit increase in TREC values estimated by the standard ddPCR corresponds to a 1.298 unit increase in TREC values estimated by Buffer 2 (Fig 3d) and this was not significantly different to 1 (95%CI for slope:0.97-1.35).

Because of the better agreement between Buffer 2 assay and the standard ddPCR than Buffer 1 assay, we took Buffer 2 forward.

#### Limit of Blank

Non template controls (NTCs) and cell line controls (CLCs) were used to estimate the LOB. None of the NTCs gave a TREC signal or a RPP30 signal (Fig 4a). Likewise, all the CLCs were negative for TRECs (Fig 4a) except for a few very high fluorescent positive droplets lying on the diagonal in a 2D plot which were artifacts and discarded in the analysis. The LOB for the crude lysate assay with buffer 2 was therefore zero.

**Figure 4:**
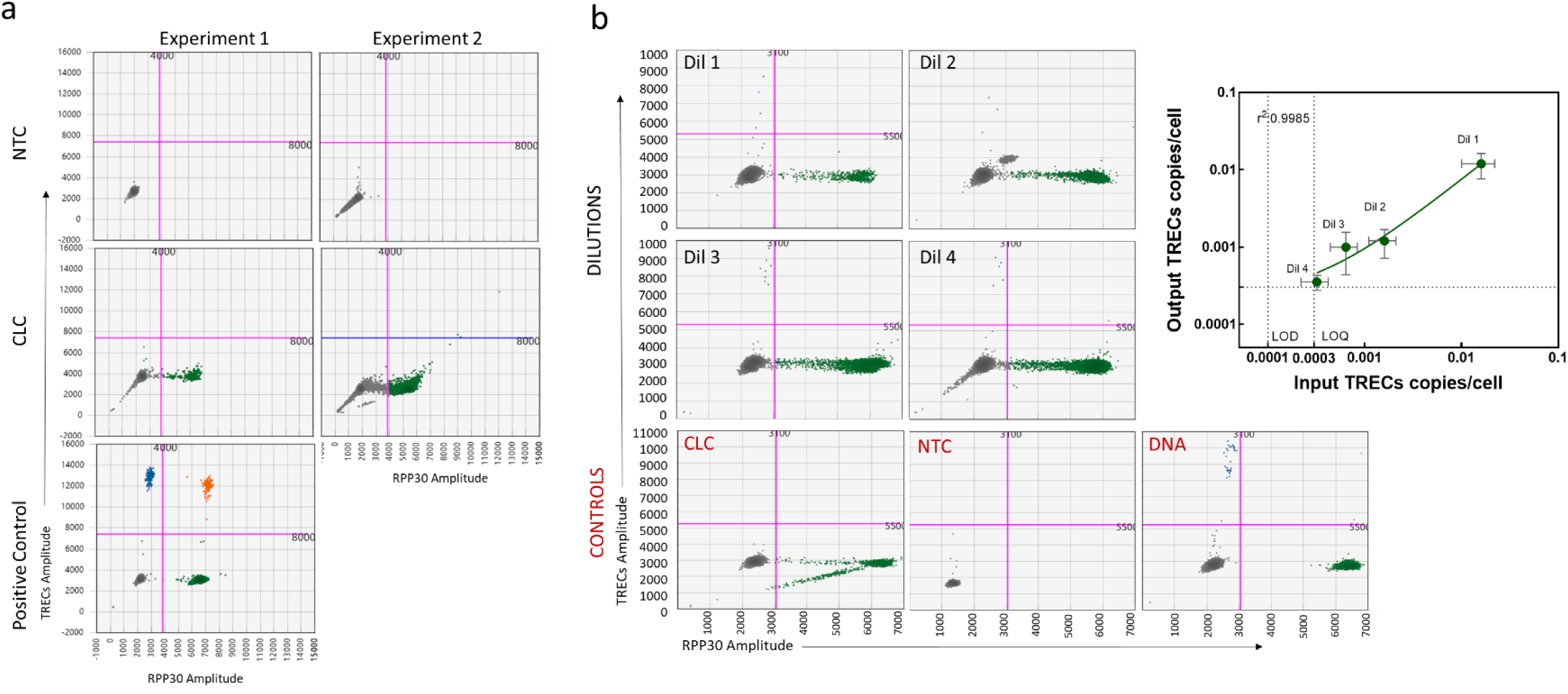
Limit of Blank (LOB), Limit of Detection (LOD) and Limit of Quantification (LOQ) of crude lysate ddPCR. (a)LOB: 2-D dot plots illustrate the results obtained from two separate LOB experiments performed on different days. The dot plots represent the merged droplets from all the samples in one run, in total, there were 22 samples for NTCs (top row) and CLCs (bottom row), 11 samples/run. In addition, a DNA sample isolated from PBMCs was run in triplicate by the standard ddPCR used as a positive control. (b) LOD and LOQ: 2-D dot plots depict the results obtained from standard curve samples made by diluting TRECs in monocyte cell line (U937) at concentrations: Dil1: 0.01 TRECs/cell, Dil2: 0.001 TRECs/cell, Dil3: 0.0004 TRECs/cell and Dil4: 0.0002 TRECs/cell. Additional controls, including CLCs, NTCs and DNA controls, are shown in the bottom three plots (labeled in red). The dot plots represent the merged droplets from triplicate dilutions. Blue dots are TRECs single positive, green dots are RPP30 single positive, orange dots are TREC RPP30 double positive and grey dots are double negative. The pink line represents the threshold. Each dot represents a droplet in ddPCR. In the right corner, linear regression analysis is shown between the average input TRECs copies/cell measured by the standard ddPCR (Dil1: 0.016±0.006 TRECs/cell, Dil2: 0.0016±0.0005 TRECs/cell, Dil3: 0.0006±0.0002 TRECs/cell and Dil4: 0.0003±0.0001 TRECs/cell) and the output TRECs copies/cell measure by crude lysate ddPCR. The coefficient of determination (r2) was calculated. Each dot represents the mean and the error bars indicate the standard deviation.

#### Limit of Detection and Quantification

To estimate the LOD of crude lysate ddPCR using Buffer 2, TREC copies were quantified from the four different TRECs concentrations made by diluting PBMCs (of known TRECs/cell, estimated by standard ddPCR), in the monocyte cell line. The concentrations of the dilutions are: Dil1: 0.01 TRECs/cell, Dil2: 0.001 TRECs/cell, Dil3: 0.0006 TRECs/cell and Dil4: 0.0003 TRECs/cell. TRECs were successfully detected in all the dilutions (Fig 4b). However, in Dil 2, an unusual cluster of double-negative appeared slightly higher diagonally than the typical double negatives, were excluded from the threshold setting manually (Fig 4b).

Since TRECs were detected in all the samples, we selected the lowest dilution, Dil4, to estimate the LOD, calculated to be 0.0001 TRECs/cell (LOD: 0+1.645*0.00007) (Fig 4b). The robustness of this detection across all dilutions was reaffirmed by 100% hit rate for probit analysis at all the concentrations. Therefore, we considered Dil4 with an average of 0.0003 TRECs/cell (95% CI 0.0002-0.0005; coefficient of variation (CV) 22%) as LOQ (Fig 4b). Finally, a linear regression between input TRECs estimated by standard ddPCR vs output TRECs estimated by ddPCR from cell lysate showed a good linearity down to 0.0003 TRECs/cells, r^2^ > 0.99, p<0.001 with good agreement between the two methods (Y = 0.7286*X + 0.0002274) (Fig 4b).

#### Intra-assay repeatability

Intra-assay repeatability was investigated by quantifying TRECs copies from the cell lysates of PBMCs (1000-8000 cells) from 4 healthy human samples. For positive control, DNA extracted from the same cells was also included in the assay. Among these four individuals, three exhibited CV% of TREC copies/reaction less than 20% (Table 1). However, one individual, LD8, displayed an outlier in one of the triplicate measurements, which resulted in a CV% above 35% (Table 1). This variation in the outlier triplicate may be attributed to the very low droplet count in this sample as compared to other triplicates, possibly due to the technical handling of the sample. On average, the CV% calculated for these four samples was 22.73% (Table 1).

**Table 1:** Intra-assay variability (Crude lysate ddPCR).

|  | TREC copies/reaction |  |  |  |  |
| --- | --- | --- | --- | --- | --- |
| Sample Id | Rep1 | Rep2 | Rep3 | AVG | CV% |
| LD8 | 20.6694 | 18.8806 | 8.8278 | 16.1259 | 39.5843 |
| TD1 | 14.9900 | 11.7225 | 13.7664 | 13.4930 | 12.2348 |
| TD2 | 33.1795 | 46.4623 | 33.7355 | 37.7924 | 19.8809 |
| DPG4 | 109.7046 | 78.9029 | 114.8296 | 101.1457 | 19.2124 |
| AVG |  |  |  |  | 22.73 |

### DIRECT COMPARISION WITH STANDARD ddPCR

Next, a direct comparison of the newly developed crude lysate ddPCR and the standard ddPCR was made by analyzing TRECs copies/cells using genuine samples (rather than T cells diluted with macrophages). The samples used were sorted naive T cells and different CD8^+^T memory subpopulations for which both the ddPCRs were run simultaneously in the same plate over different days for different samples. Mean TREC copies/cell were not significantly different between the two assays (Wilcoxon test, p = 0.31) (Fig 5a). The Bland-Altman analysis indicated a bias of -0.001635 between two methods with SD of 0.008337, which indicates that, on average, standard ddPCR tends to slightly underestimate the TREC copies compared to ddPCR directly with cell lysate (Fig 5b). The small standard deviation suggests the differences between the methods are relatively consistent and that the level of agreement between the methods is relatively high.

**Fig 5:**
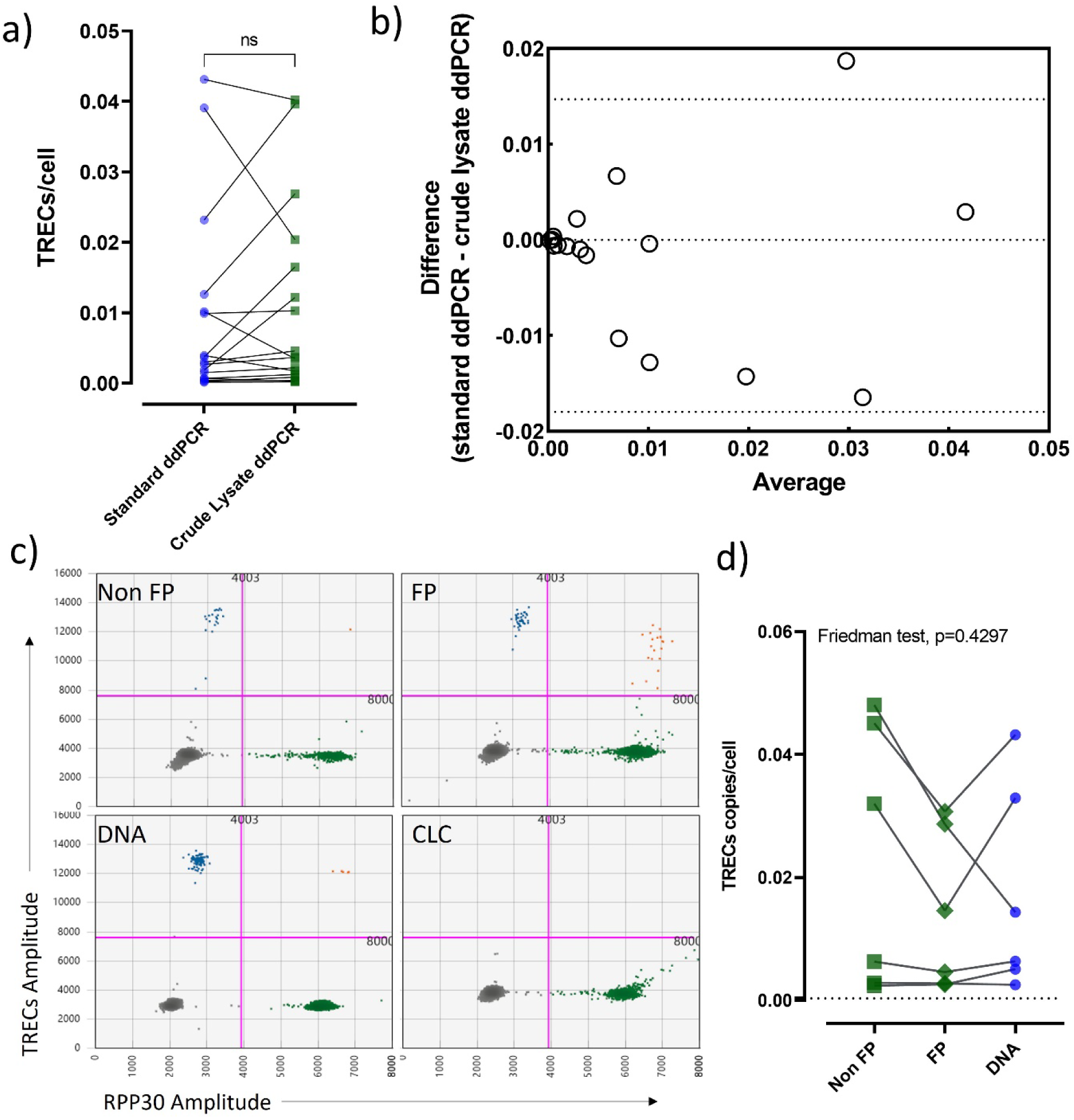
Accuracy of crude lysate ddPCR and its utility on fixed and permeabilized (FP) cells. (a) A pairwise comparison between TRECs copies/cell detected by the standard ddPCR (in blue circles) and crude lysate ddPCR (in green squares) is plotted. Each symbol represents a donor, and pairwise comparison is shown by connecting black lines. Stats: Wilcoxon test, ns=non significant (p>0.05). (b) The Bland-Altman analysis is plotted as a scatter diagram with the TREC copies/cell differences between the two methods on the y-axis and the averages of these two measurements on the x-axis. Horizontal lines are drawn at the mean difference and at the limits of agreement. (c) The 2-D dot plots display TRECs and/or RPP30 positive droplets detected from paired FP, non FP and extracted DNA samples, representation of only one sample. Additionally, CLC, a negative control, is also shown. In the dot plots, blue dots are TRECs single positive, green dots are RPP30 single positive, orange dots are TREC RPP30 double positive and grey dots are double negative. The pink line represents the threshold. Each dot represents a droplet in ddPCR.(d) A pairwise comparison of TRECs copies/cells obtained from FP (in green diamonds), non FP (in green squares) and extracted DNA (in blue circles) samples was conducted. Each symbol represents a donor, and pairwise comparison is shown by connecting black lines. Stats: Friedman test with Dunn’s multiple comparisons, denoting significance as *<0.05.

### IMPACT OF CELL FIXATION AND PERMABILIZATION

Extracting DNA from fixed and permeabilized cells is challenging due to the crosslinking of proteins and nucleic acids caused by the fixing process. However, it is often necessary to work with fixed, permeabilized samples. We therefore investigated the impact of fixation and permeabilization on TRECs quantification using crude lysate ddPCR using buffer 2. Six different samples of fixed and permeabilized cells were compared with paired non-fixed cells and extracted DNA (standard ddPCR) samples (Fig 5 c). Results for TRECs count were not significantly different between these three different groups (p=0.4) (fixed (mean=0.014), non-fixed cells (mean=0.022) and standard ddPCR (mean=0.017)). However the TREC count tended to be lower in fixed cells compared to non-fixed cells for the three samples with a higher TREC concentration (Fig 5d). We suggest that additional optimization is conducted before using this assay on fixed and permeabilized cells.

## Discussion

Accurate quantification of rare genes from limited clinical samples is challenging yet crucial for research purposes, as it is used in the diagnosis of rare diseases, for investigating disease mechanisms, for biomarker identification, and for assessing treatment response. However, traditional nucleic acid extraction methods may lead to target loss. Therefore, it is necessary to develop alternative protocols. To address this, in the present study, various methods including thermal lysis, sonication, lysis buffer from DNA extraction kit, Ambion Cell to-Ct® kit (Buffer 1) and SuperScript™ IV CellsDirect™ cDNA Synthesis Kit (Buffer 2) were tested to make the cell lysate compatible with ddPCR to bypass the DNA extraction step. We also introduced a viscosity breakdown step that overcame the increased viscosity of crude lysates (Fig 2) which emerged as a critical challenge in this process. Among the methods evaluated, the combination of buffer 1 or 2 with the viscosity breakdown step exhibited promising results (Fig 4), with buffer 2 emerging as the preferred choice.

Unexpectedly we found that the droplet volume obtained using cell lysate (0.70nL) was lower than that assumed by QuantaSoft v1.0.569 (0.85nL). This decrease in volume from that expected was not attributable to the cell lysate protocol as we found an almost identical droplet volume with the standard ddPCR assay (0.70nl) (Fig 3). This observation is consistent with previous studies that have also shown deviations from the default droplet volume in Biorad’s system (17,20,21). Notably, Bogozalec Kosir et al. (2017) reported a droplet volume of 0.71nL, very similar to our findings (21).

The accuracy of this protocol was critically assessed and compared to the standard ddPCR protocol. Overall, the novel assay showed good agreement with the standard assay in the range from 0.0003 to 0.01 TRECs/cell. This was confirmed with populations isolated from PBMC (a more representative scenario).

Quantifying the performance of the new assay we found the LOB to be zero, the LOD to be <0.0001 TREC/cell and the LOQ to be <0.0003 TRECs/cell. This is favourable compared to standard ddPCR for which we found the LOB to be zero, the LOD to be 0.0004 TRECs/ cell and the LOQ to be 0.002 TRECs/cell. This relatively high LOQ for the standard assay is consistent with previous reports. Profaizer & Slev (2020) determined the LOQ for TRECs using standard ddPCR to be 2 copies/μl of blood (23) and for qRT-PCR to be 24 copies/μl of blood (23). To compare these results with our study, we transformed the units by estimating the number of PBMCs per μl of blood (1 μl of blood yields 1*10^3^ - 2*10^3^ PBMCs), this translates to a LOQ of 0.002 - 0.001 TRECs/cell for the standard ddPCR and 0.024 - 0.012 TRECs/cell for qRT-PCR, approximately 3 and 40 times higher, respectively, than our novel assay.

The average CV% for the repeatability experiment is 22.72% suggesting some source of variability (Table 2). The slight variability observed in these samples may arise from the viscosity of PCR mix, droplet count variations, incomplete mixing of samples or the presence of secondary DNA structures, which might disturb the random distribution of samples. Furthermore, variations in sample handling, pipetting or minor inconsistencies in assay conditions could also impact the assay’s consistency. Further investigations, such as fine-tuning by PCR enhancers or cycling conditions, may offer the potential to enhance assay consistency further.

By partitioning a single sample of cell lysates into four ddPCR reactions (to fully consume the lysate), we were able to comprehensively screen the entire sample to detect rare genes. Subsequently, these four samples were merged to calculate absolute quantification. This approach significantly increased the number of screened droplets per sample, from about 20,000 per reaction or sample to 80,000 per sample.

Our method is preferable to those using a pre-amplification reaction. Pre-amplification involves increasing the copy number of nucleotide sequences in the reaction before polymerase chain reaction (PCR) analysis, enabling the analysis of genes with limited nucleic acid content. In pre-amplification reactions, addressing two major challenges is crucial: augmenting the reaction’s capacity and ensuring target amplification specificity. This ensures efficient amplification during each PCR cycle, even for targets with significantly different initial quantities and substantial background noise (25). Failing to tackle these challenges can introduce significant bias. By circumventing the need for pre-amplification, we eliminated related issues. Simply put, fewer steps in the process reduce the potential for introducing variability.

Our method differs from previously published studies on crude lysate ddPCR for absolute quantification of genes (26–28), as we utilized the whole cell lysate instead of a portion of it. Working with a larger volume of cell lysate presents challenges as it alters the viscosity of the reaction mixture, resulting in a reduction in droplet count and experiment efficiency. However, in our study, we successfully addressed this challenge by introducing a viscosity breakdown step. This approach is more robust for detecting rare events from limited clinical or biological samples.

Furthermore, we conducted experiments to evaluate the efficacy of this protocol on fixed cells. Although there were no significant differences observed between fixed/permeabilized cells and non-fixed cells, it was noted that the count of TRECs tended to be lower in the fixed cells compared to the non-fixed cells. We suggest that further optimization is needed before this method can be used on fixed and permeabilized samples.

It has been established that using plasmids for optimizing qPCR reactions is unsuitable as it can lead to overestimation of the target ((Hou et al. 2010). Therefore, for the crude lysate ddPCR optimisation, we used PBMCs rather than plasmids containing TRECs DNA. In adherence to MIQE guidelines, we consistently included appropriate negative and positive controls in our experiments to assess background signals and set the threshold (24).

The study’s limitations include a relatively small number of replicates used for optimization and technical assessment. Furthermore, while our LOD and LOQ assessment indicated the presence of TRECs copies in all dilutions, we did not explore concentrations below 0.0003 TRECs copies/cell due to the satisfactory performance within the optimized range so these represent upper bounds. We encountered occasional artifacts, such as double negative clusters (Figure 3.6 Dil2) or isolated droplets with significantly higher TRECs and RPP30 amplitudes than the rest. These artifacts may be attributed to contamination, debris in the crude lysate, or technical variations, such as temperature fluctuations or minor inconsistencies during droplet generation.

In summary, the novel protocol of crude lysate ddPCR is accurate, sensitive, and precise in estimating rare gene from small samples. By eliminating the DNA extraction step and utilizing the whole cell lysate, the protocol minimizes potential sources of variability, such as DNA loss, and provides a robust approach for the accurate analysis of rare gene events.

## Ethics Statement

All study procedures were conducted according to the principles of the Declaration of Helsinki and all participants gave written informed consent. Human samples used in this research project were obtained from the Imperial College Healthcare Tissue and Biobank (ICHTB). ICHTB is supported by the National Institute for Health Research (NIHR) Biomedical Research Centre based at Imperial College Healthcare NHS Trust and Imperial College London. ICHTB is approved by Wales REC3 to release human material for research (22/WA/0214).

## Acknowledgements

The authors would like to thanks Dr. Danai Koftori for taking blood and wish to acknowledge the support of the St Mary’s Flow Cytometry Core Facility at Imperial. This research was funded by European Union’s Horizon 2020 research and innovation programme grant 764698 (QUANTII).

## Author contributions

CK performed the experiments, performed the data analysis and wrote the manuscript. SA performed the experiments. CNK supervised the project and helped write the manuscript. BA conceived the project, obtained funding, supervised the project and helped write the manuscript.

## References

1. Taylor SC, Laperriere G, Germain H. Droplet Digital PCR versus qPCR for gene expression analysis with low abundant targets: From variable nonsense to publication quality data. Sci Rep. 2017;7(1).

2. Pharo HD, Andresen K, Berg KCG, Lothe RA, Jeanmougin M, Lind GE. A robust internal control for high-precision DNA methylation analyses by droplet digital PCR. Clin Epigenetics. 2018;10(1).

3. Levy CN, Hughes SM, Roychoudhury P, Reeves DB, Amstuz C, Zhu H, et al. A highly multiplexed droplet digital PCR assay to measure the intact HIV-1 proviral reservoir. Cell Rep Med. 2021;2(4).

4. Bell AD, Usher CL, McCarroll SA. Analyzing copy number variation with droplet digital PCR. In: Methods in Molecular Biology. 2018.

5. Lu A, Liu H, Shi R, Cai Y, Ma J, Shao L, et al. Application of droplet digital PCR for the detection of vector copy number in clinical CAR/TCR T cell products. J Transl Med. 2020;18(1).

6. Rowlands V, Rutkowski AJ, Meuser E, Carr TH, Harrington EA, Barrett JC. Optimisation of robust singleplex and multiplex droplet digital PCR assays for high confidence mutation detection in circulating tumour DNA. Sci Rep. 2019;9(1).

7. Roberts C h., Jiang W, Jayaraman J, Trowsdale J, Holland MJ, Traherne JA. Killer-cell Immunoglobulin-like Receptor gene linkage and copy number variation analysis by droplet digital PCR. Genome Med. 2014;6(3).

8. Heredia NJ. Droplet digital ^TM^ PCR next-generation sequencing library QC assay. In: Methods in Molecular Biology. 2018.

9. Gutiérrez-Aguirre I, Rački N, Dreo T, Ravnikar M. Droplet digital PCR for absolute quantification of pathogens. Methods in Molecular Biology. 2015;1302.

10. Hindson BJ, Ness KD, Masquelier DA, Belgrader P, Heredia NJ, Makarewicz AJ, et al. High-throughput droplet digital PCR system for absolute quantitation of DNA copy number. Anal Chem. 2011;83(22).

11. Serana F, Chiarini M, Zanotti C, Sottini A, Bertoli D, Bosio A, et al. Use of V(D)J recombination excision circles to identify T- and B-cell defects and to monitor the treatment in primary and acquired immunodeficiencies. Vol. 11, Journal of Translational Medicine. 2013.

12. Bausch-Jurken MT, Verbsky JW, Routes JM. Newborn screening for severe combined immunodeficiency - A history of the TREC assay. Vol. 3, International Journal of Neonatal Screening. 2017.

13. Gattinoni L, Lugli E, Ji Y, Pos Z, Paulos CM, Quigley MF, et al. A human memory T cell subset with stem cell-like properties. Nat Med. 2011;17(10).

14. Lugli E, Gattinoni L, Roberto A, Mavilio D, Price DA, Restifo NP, et al. Identification, isolation and in vitro expansion of human and nonhuman primate T stem cell memory cells. Nat Protoc. 2013;8(1).

15. Bacher P, Scheffold A. Flow-cytometric analysis of rare antigen-specific T cells. Cytometry Part A. 2013;83(8).

16. Sandgaard KS, Lewis J, Adams S, Klein N, Callard R. Antiretroviral therapy increases thymic output in children with HIV. AIDS. 2014;28(2).

17. Pinheiro LB, Coleman VA, Hindson CM, Herrmann J, Hindson BJ, Bhat S, et al. Evaluation of a droplet digital polymerase chain reaction format for DNA copy number quantification. Anal Chem. 2012;84(2).

18. Milosevic D, Mills JR, Campion MB, Vidal-Folch N, Voss JS, Halling KC, et al. Applying standard clinical chemistry assay validation to droplet digital PCR quantitative liquid biopsy testing. Clin Chem. 2018;64(12).

19. Menacho-Melgar R, Lynch MD. Measuring Oligonucleotide Hydrolysis in Cellular Lysates via Viscosity Measurements. Bio Protoc. 2022;12(2).

20. Dong L, Meng Y, Sui Z, Wang J, Wu L, Fu B. Comparison of four digital PCR platforms for accurate quantification of DNA copy number of a certified plasmid DNA reference material. Sci Rep. 2015;5.

21. Košir AB, Divieto C, Pavšič J, Pavarelli S, Dobnik D, Dreo T, et al. Droplet volume variability as a critical factor for accuracy of absolute quantification using droplet digital PCR. Anal Bioanal Chem. 2017;409(28).

22. Tessitore MV, Sottini A, Roccaro AM, Ghidini C, Bernardi S, Martellosio G, et al. Detection of newly produced T and B lymphocytes by digital PCR in blood stored dry on nylon flocked swabs. J Transl Med. 2017;15(1).

23. Profaizer T, Slev P. A multiplex, droplet digital PCR assay for the detection of T-cell receptor excision circles and kappa-deleting recombination excision circles. Clin Chem. 2020;66(1).

24. Whale AS, De Spiegelaere W, Trypsteen W, Nour AA, Bae YK, Benes V, et al. The Digital MIQE Guidelines Update: Minimum Information for Publication of Quantitative Digital PCR Experiments for 2020. Clin Chem. 2020;66(8).

25. Okino ST, Kong M, Sarras H, Wang Y. Evaluation of bias associated with high-multiplex, target-specific pre-amplification. Biomol Detect Quantif. 2016;6.

26. Ludlow AT, Shelton D, Wright WE, Shay JW. DdTRAP: A method for sensitive and precise quantification of telomerase activity. In: Methods in Molecular Biology. 2018.

27. Zou Z, Guo L, Ahmadi P, Hartjen P, Gosau M, Smeets R, et al. Two simple and inexpensive methods for preparing DNA suitable for digital PCR from a small number of cells in 96-well plates. J Clin Lab Anal. 2021;35(1).

28. Vasudevan HN, Xu P, Servellita V, Miller S, Liu L, Gopez A, et al. Digital droplet PCR accurately quantifies SARS-CoV-2 viral load from crude lysate without nucleic acid purification. Sci Rep. 2021;11(1).

